# WDR5 and Myc Cooperate to Regulate Formation of Neural Crest Stem Cells

**DOI:** 10.1101/2025.09.19.677424

**Authors:** Karlin Compton, Elizabeth Barter, Carole LaBonne

## Abstract

WDR5 is a multifunctional scaffolding protein with established roles in chromatin regulation and pluripotency, but its functions in early vertebrate development remain poorly understood. Here, we show that *Xenopus wdr5* is expressed in blastula stem cells and enriched in neural crest cells during neurulation. Morpholino-mediated depletion of wdr5 abolished neural crest marker expression both in embryos and in reprogrammed explants while unexpectedly expanding neural plate border and neural plate domains. Gain-of-function experiments revealed a striking dose-dependent effect: low levels of wdr5 enhanced neural crest formation, whereas high levels suppressed it, suggesting a requirement for precise stoichiometry with interacting partners. We identify myc as a critical cofactor for wdr5 in neural crest specification—wdr5 and myc physically interact in early embryos, and co-expression at defined ratios rescues neural crest formation while either, individually, is inhibitory. Domain-specific mutagenesis showed that the WBM site of wdr5 is required for myc-dependent activation of neural crest genes, while the WIN site regulates *myc* expression itself; both domains are necessary to rescue wdr5 loss-of-function phenotypes. Moreover, modulation of myc levels phenocopied wdr5’s dose-sensitive effects, reinforcing the importance of balanced wdr5–myc activity. These findings reveal that wdr5 orchestrates neural crest development through multiple, domain-specific mechanisms—integrating stoichiometric control with partner-specific transcriptional regulation—and highlight parallels with Wdr5–Myc cooperation in cancer, underscoring the broader relevance of precise cofactor ratios in cell fate decisions.

## Introduction

The neural crest, a stem cell population unique to vertebrates, contributes a large and diverse set of cell types to the vertebrate body plan including much of the peripheral nervous system, melanocytes, and craniofacial bone and cartilage (Le Douarin & Kalcheim, 1999). Because acquisition of these cells drove the evolution of vertebrates, a deeper understanding of the mechanisms regulating formation of neural crest cells will provide key insights into vertebrate origins, and how these cells contribute such a wide range of specialized cell types to the vertebrate body plan (Schock et al., 2023).

Neural crest cells arise in the ectoderm at the neural plate border and retain multi-germ layer developmental potential past when most embryonic cells have become lineage restricted (Prasad et al., 2012). Insights into the origins of neural crest potential came from the realization that these stem cells share gene regulatory network (GRN) components, and other features, with pluripotent stem cells of the vertebrate blastula (Buitrago-Delgado et al., 2015; York et al., 2024) Both pluripotent blastula and neural crest stem cells express core neural crest regulatory genes such as *snai1*, *id3*, *foxd3*, *tfap2a* as well as core pluripotency factors *myc*, *ventx/nanog*, *pou5* and *sox2/3* (Bellmeyer et al., 2003; Buitrago-Delgado et al., 2015; York et al., 2024). These two stem cell populations share additional features including high levels of Erk activity, low levels of histone acetylation and a requirement for BET family epigenetic readers (Geary & LaBonne, 2018; Huber et al., 2024; Rao & LaBonne, 2018). HDAC1 can enhance the generation of neural crest cells and has also been shown to cooperate with NANOG to promote pluripotency in murine and human embryonic stem cells (Bogdanovic et al., 2012; Dovey et al., 2010; Rao & LaBonne, 2018; Watanabe et al., 2013).

To gain further insights into the control of developmental potential in neural crest stem cells, we sought to identify new regulators of these stem cell populations. Using previously published datasets of the transcriptome changes that occur when blastula stem cell explants are reprogrammed to a neural crest state (Huber and LaBonne, 2024), we identified *wdr5* (*WD40 repeat domain containing protein 5*) as a factor significantly upregulated in response to neural crest induction. wdr5 is a highly conserved protein that shares more than 90% sequence identity across vertebrates (Schuetz et al., 2006). It is known for scaffolding histone methyltransferase complexes (COMPASS) as well as for interacting with diverse transcription factors (Guarnaccia & Tansey, 2018). *wdr5* is robustly expressed in mouse embryonic stem cells (mESCs) and its expression decreases as these cells differentiate (Ang et al., 2011). wdr5 is involved in the formation of a number of histone modifying complexes in stem cells, and may also play a role in reading histone modifications (Guarnaccia & Tansey, 2018; Ruthenburg et al., 2006; Schuetz et al., 2006; Wysocka et al., 2005).

Recent evidence indicates that wdr5’s role in depositing H3K4me3 explains only a subset of its effects on transcription, whereas others may be mediated through interactions with DNA-binding transcription factors (Siladi et al., 2022). wdr5 has been shown to interact with a number of transcription factors including p53, Twist, Pou5f1/Oct4 and Myc (Ang et al., 2011). Physical interactions with wdr5 are largely mediated by one of two highly conserved binding sites on opposite sides of beta-propeller structure of wdr5: the wdr5 binding motif (WBM), or the wdr5 interaction domain (WIN) (Dharmarajan et al., 2012; Guarnaccia & Tansey, 2018; Thomas, Wang, et al., 2015).

In this study, we investigate the role of wdr5 in neural crest stem cells in *Xenopus*. We show that *wdr5* is required for formation of the neural crest in whole embryos and that depletion of *wdr5* prevents pluripotent blastula explants from adopting a neural crest fate. Additionally, we show that exogenous wdr5 expression alters neural crest factor expression in a dose-dependent manner, with higher concentrations exhibiting an inhibitory effect on neural crest gene expression, suggesting stoichiometric effects. We show that *wdr5* and the pluripotency/neural crest transcription factor myc physically interact with one another in *Xenopus* and colocalize to the same cells at neural crest stages. Co-expression of *wdr5* and *myc* together facilitates an expansion of the neural crest domain. Finally, we show find that mutations in either of the two conserved interaction domains alters the ability of wdr5 to regulate neural crest formation in distinct ways, and that binding between *wdr5* and *myc* is required for the regulation of neural crest formation.

## Results

### wdr5 is expressed in blastula stem cells and is upregulated in neural crest cells

As a first step in studying the role of wdr5 in early embryonic *Xenopus* development we characterized its expression. Analysis of previously published RNA-Seq data (Huber and LaBonne, 2024) showed that *wdr5* is highly expressed in blastula stems cells, at levels comparable to factors involved in pluripotency including *ventx2.2* and *tfap2a*. Wnt/Chordin mediated-reprogramming of animal pole explants to a neural crest state promoted expression of *wdr5* at both early and late neurula stages (Fig 1A). We used whole mount *in situ* hybridization (WISH) to examine the spatial expression of *wdr5*. Consistent with the RNA-Seq data, *wdr5* is strongly expressed in the pluripotent animal pole cells of blastula stage embryos (Fig. 1B). At early and mid-neurula stages *wdr5* expression is enriched in the developing central nervous system (CNS) and neural crest cells, and at tailbud stages, *wdr5* is enriched in migratory neural crest cells. We used fluorescent *in situ* Hybridization Chain Reaction (HCR-FISH) to examine if *wdr5* is co-expressed with neural crest factors. We observed that *wdr5* is co-expressed with *myc* in the neural crest at neurula stages (Fig. 1C). Taken together, the spatial and temporal expression profile of *wdr5* is consistent with a role in regulating neural crest development.

**Figure 1.**
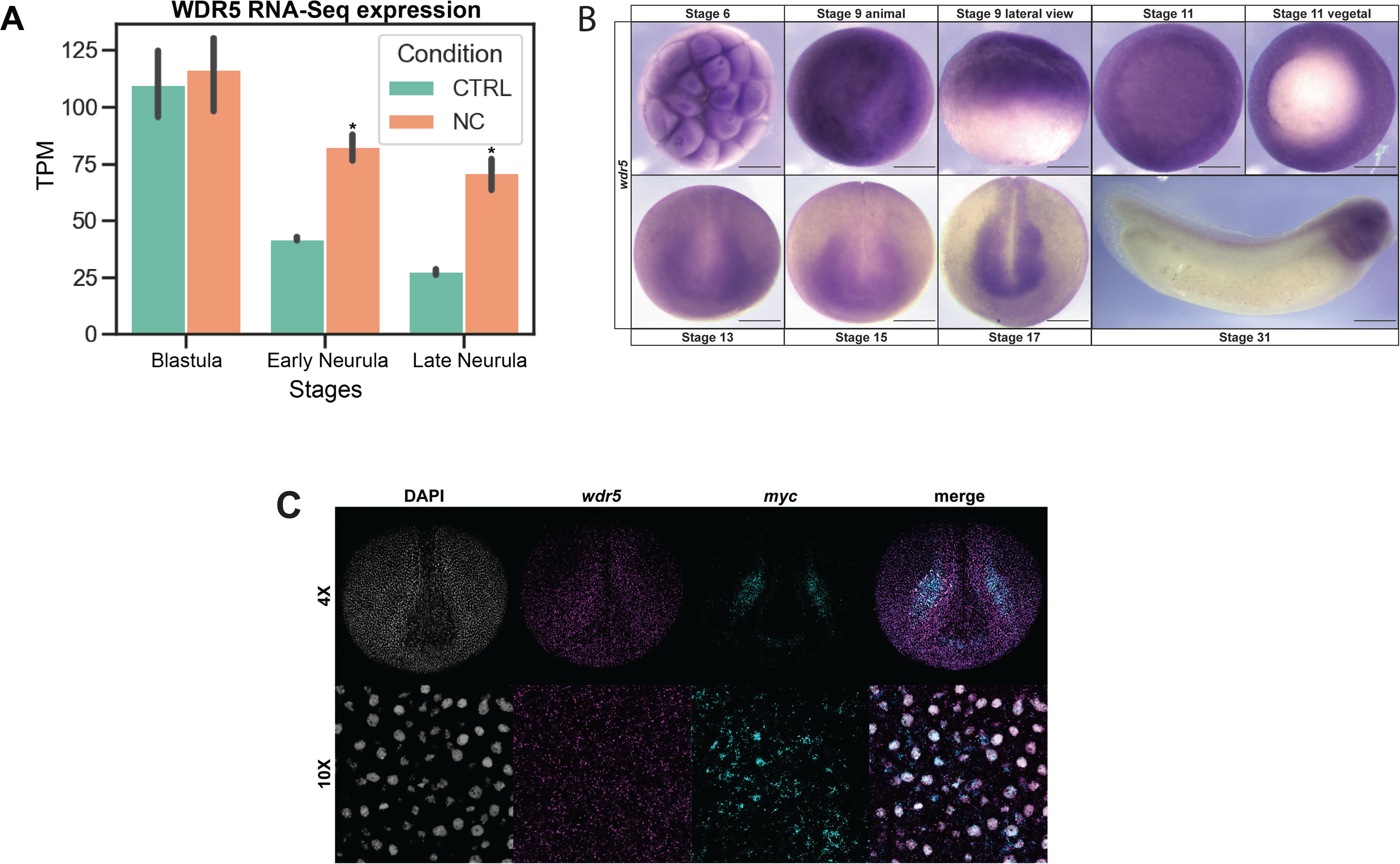
Characterization of *wdr5* expression. (A) Analysis of previously published RNA-Seq data (York et al., 2024) shows *wdr5* transcripts are abundant in the animal pole cells of the blastula stages and differentially enriched in neural crest-induced animal caps at early and late neurula stages. (p < 0.05) (B) Spatial and temporal expression of *wdr5* shows that *wdr5* transcripts are maternally provided, enriched in blastula stem cells and retained broadly throughout the neuroectoderm into developing neural crest stem cells and neural crest derivatives. (C) HCR-FISH confirms *wdr5* expression in the neural crest that co-localizes with myc – a canonical neural crest and pluripotency factor. Abbreviations: [[CTRL = control, NC = neural crest]]

### wdr5 is essential for neural crest formation and patterning of the neural plate border

After finding that *wdr5* expression is enriched in neural crest cells, we next asked whether it is essential for neural crest formation. To test this, we designed a *wdr5* translation-blocking morpholino oligonucleotide (MO) that targets both allo-alleles of *wdr5*. Co-injection of the MO with mRNA encoding a c-terminally myc-tagged wdr5 led to loss of *wdr5* expression as assayed by western blot (Supp Fig. 1A). We performed targeted injection into 2 animal pole blastomeres on one side of 8-cell *Xenopus* embryos to target the neural crest, with beta-galactosidase co-injected as a lineage tracer. Embryos were cultured to neurula stages and examined by WISH for expression of neural crest markers. Depletion of *wdr5* resulted in a loss of expression of neural crest factors *foxd3* (86%, *n*=74) and *snai2* (94%, *n*=34), indicating that wdr5 activity is essential for neural crest formation (Fig 2A). Neural crest could be rescued by co-injecting mRNA encoding n-terminally Flag tagged *wdr5* (*foxd3 (*70% rescued, n=54); *snai2* (72% rescued, n=39) compared to the loss of neural crest factor expression in embryos injected with only wdr5 MO (*foxd3*: 92% loss, n=60; *snai2*: 82% loss, n=35) (Fig. 2B,C).

**Figure 2.**
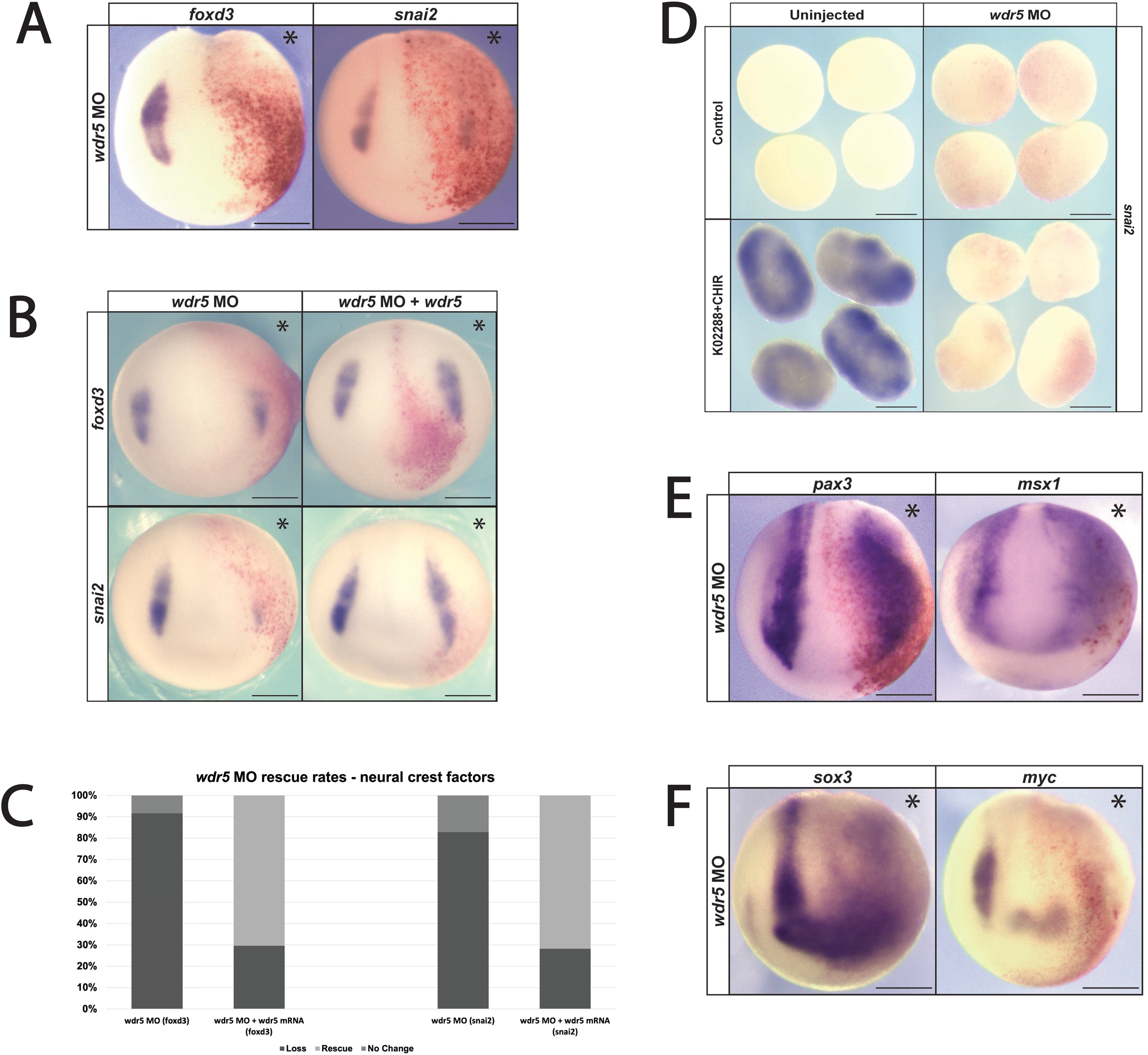
Morpholino-mediated knockdown shows that *wdr5* is required for NC gene expression. (A) Morpholino-mediated depletion of *wdr5* inhibits neural crest factor expression. (B) Expression of *wdr5* mRNA rescues morpholino-mediated neural crest gene expression. (C) Quantification of rescue percentage. (D) Morpholino-mediated depletion of *wdr5* prevents pluripotent blastula cells from being reprogrammed to neural crest state. (E) Morpholino-mediated depletion of *wdr5* does not inhibit expression of neural plate border and neural plate factors. (F) Knockdown of *wdr5* leads to an expansion of the neural plate factor *sox3* and loss of *myc* expression at neurula stages. Abbreviations [[MO = morpholino]]

To determine if wdr5 is also required for reprogramming pluripotent blastula cells to a neural crest state we treated explants from control or morpholino-injected embryos with a small molecule inhibitor of BMP signaling (k02288) and an agonist of Wnt signaling (CHIR) to induce neural crest (Huber & LaBonne, 2024). Morpholino-mediated depletion of *wdr5* prevented reprogramming to a neural crest state, as evidenced by a failure to induce the expression of *snai2* (100%, *n*=54) compared to controls (Fig. 2D).

### Loss of wdr5 enhances neural plate border formation

We next asked if *wdr5* is specifically required for neural crest gene expression or alternatively is also essential for establishing the neural plate border (NPB) region. The NPB is characterized by the overlapping expression of several transcription factors, including *pax3, zic1* and *msx1 (*Groves and LaBonne, 2014). Surprisingly, we found that depletion of wdr5 led to expanded expression of the NPB factors *pax3* (85% expansion, *n*=47) and *msx1* (82% expansion, *n*=46), as well as neural plate marker *sox3* (93% expansion, *n*=42), into regions that normally form neural crest and epidermis (Fig 2E,F). Thus, *wdr5* is not required for formation of all ectoderm-derived cell types. Interestingly, expression of *myc* at the NPB did require wdr5 (90% loss, n=54).

### Morpholino-mediated wdr5 depletion does not significantly impact global H3K4me3 levels

Wdr5 can serve as a scaffolding protein for MLL/SET methyltransferases, which facilitate trimethylation of Histone 3 at lysine 4 (H3K4me3), an epigenetic modification associated with transcriptional activation (Bernstein et al., 2002; Dou et al., 2006). We therefore examined the effects of *wdr5* depletion on global methylation levels by comparing H3K4me3 levels in control explants with explants depleted for *wdr5*. Control embryos or embryos injected with wdr5 morpholino in both cells at the 2-cell stage were cultured to blastula stages when animal caps were excised and cultured to neurula stages. Western blot analysis showed that wdr5-depleted explants and control explanted exhibited comparable levels of H3K4me3 as normalized to total histone H3 (Supp. Fig. 1B). Thus, the loss of neural crest cells in wdr5-depleted embryos is independent of major changes in global methylation levels.

### wdr5 activity affects neural crest formation in a dose-dependent manner

Given that wdr5 is required for neural crest formation we next asked whether increasing wdr5 levels might promote neural crest formation. mRNA encoding wdr5 was injected unilaterally into two animal pole cells of four-cell *Xenopus* embryos targeting the presumptive neural crest, with the uninjected side serving as an internal control. Injections were carried out using a range of mRNA concentrations to identify potential dose-dependent effects. Lower levels of wdr5 were found to enhance the expression of neural crest factors *foxd3* (70%, n=54) and *snai2* (75%, n=48) and neural plate boarder markers *pax3* (90%, *n*=83) and *msx1* (87%, *n*=62) (Fig. 3A,B). However, whereas higher levels of wdr5 similarly enhanced neural plate boarder gene expression (*pax3*: 93.75% expansion, n=48, *msx1*: 90% expansion, n=40) those same doses inhibited expression of neural crest markers (*foxd3*, 80%, *n*=49; *snai2*, 74%, *n*=57) (Fig 3A,B). The striking concentration-dependent effects of wdr5 on neural crest factors suggests that its function is likely dependent on interaction partners sensitive to stoichiometry. By contrast, the observed enhancement of neural plate boarder gene expression by both high and low levels of wdr5 indicates that wdr5 functions somewhat differently in the regulation of these two cell types (Supp Fig. 2A).

**Figure 3.**
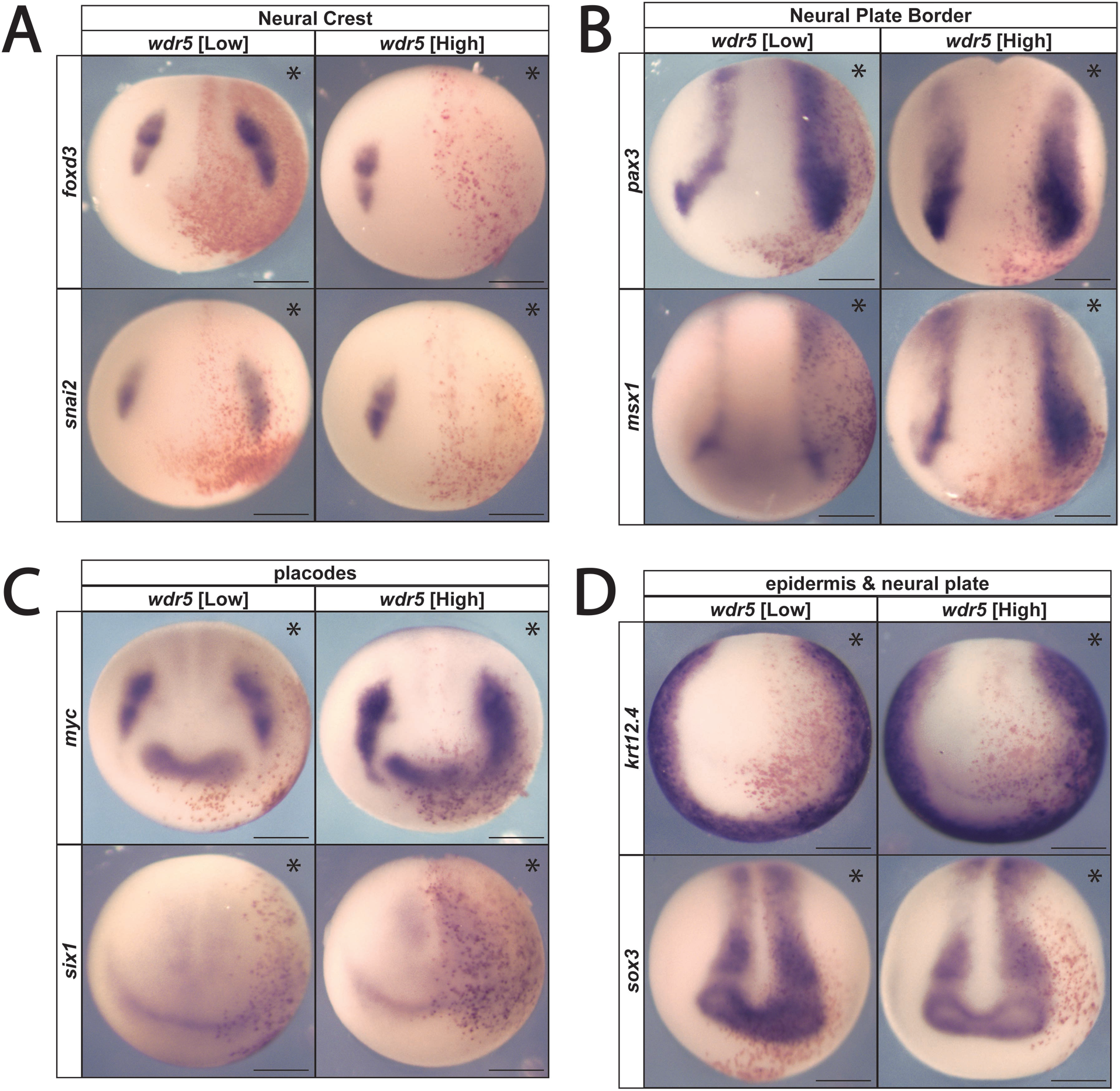
Overexpression of *wdr5* exhibits concentration-dependent effects on neural crest factor expression. (A) Exogenous *wdr5* mRNA shows concentration dependent effects on neural crest gene expression. (B) Increased *wdr5* expression expands neural plate border gene expression independent of mRNA concentration. (C) Exogenous *wdr5* mRNA exhibits expanded expression of *myc* expression and inhibition of *six1* expression. (D) Increased *wdr5* expression inhibits expression of epidermal keratin factor *krt12.4* and expansion of the neural plate factor *sox3* independent of mRNA concentration.

### wdr5 and Myc physically interact

We have previously shown that the proto-oncogene myc is required for neural crest formation (Bellmeyer et al., 2003; Light et al., 2005). Myc is also a known wdr5-interacting factor that binds to its WBM site (Fig. 4A). Wdr5 has been shown to interact directly with Myc in human cancer stem cells, and is required for myc-mediated transcriptional regulation (Thomas, Wang, et al., 2015; Thomas et al., 2019; Thomas, Foshage, et al., 2015). Accordingly, we asked if wdr5 and myc can physically interact in Xenopus. mRNA encoding flag-tagged wdr5 and myc-tagged myc was injected into 1 cell of 2-cell *Xenopus* embryos, embryos were lysed at blastula stages, and myc was immunoprecipitated from lysates using an anti-myc antibody. Western blot analysis using an anti-flag antibody showed a robust physical interaction between wdr5 and myc (Fig 4B).

**Figure 4.**
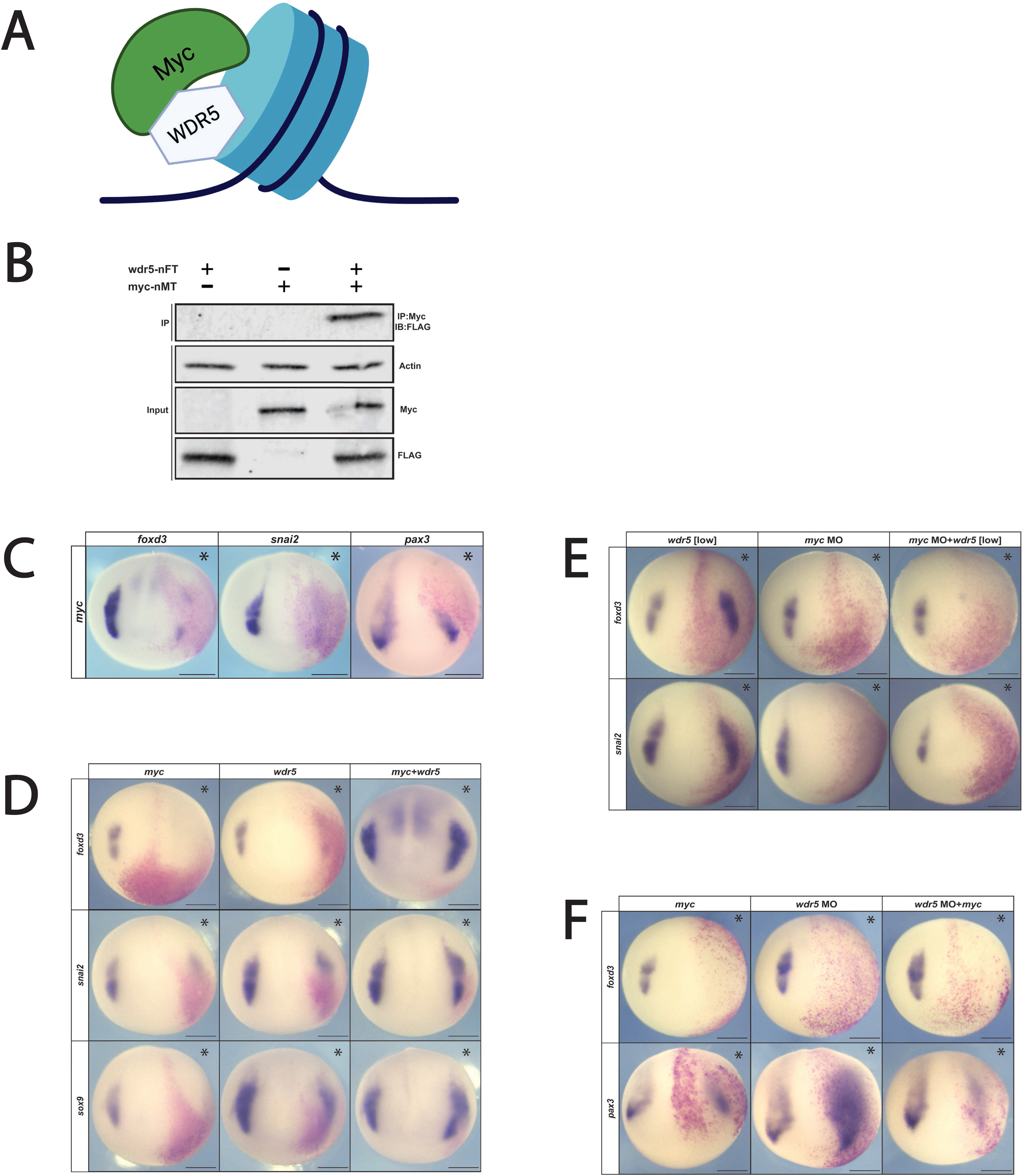
*myc* and *wdr5* interact directly to facilitate neural crest factor expression. **(A)** *wdr5* and *myc* directly interact at chromatin to influence downstream gene expression of myc target genes. Western blot of Co-IP shows that *wdr5*-FLAG and *myc-*Myc directly interact in *Xenopus* embryos. *myc* mRNA overexpression inhibits neural crest and neural plate border gene expression. (D) Co-expression of *wdr5* mRNA and *myc* mRNA facilitates neural crest gene expression. (E) *myc* expression is required for wdr5-mediated neural crest expansion. (F) *myc* mRNA inhibits neural crest and neural plate border gene expression when *wdr5* expression is downregulated.

### Myc overexpression inhibits neural crest formation

As a first step toward examining if there is a joint role for wdr5 and myc in neural crest formation we examined the effects of myc up-regulation on neural crest and neural plate border formation. mRNA encoding myc was injected at a range of concentrations into 2 blastomeres of 4-cell *Xenopus* embryos targeting the presumptive neural crest. We found that at all concentrations myc inhibited expression of neural crest markers *foxd3* (83.3% loss, n=48) and *snai2* (77.7% loss, n=45) (Fig 4C and not shown). Surprisingly, myc-upregulation also reduced expression of the neural plate border factor *pax3* (Fig. 4C) (89.7% loss, n=29), in contrast to what was observed for wdr5 gain-of-function. This finding is consistent with myc having both wdr5-dependent and -independent functions.

### Co-expression of wdr5 and myc promotes neural crest formation

If the relative expression levels of Myc and wdr5 are important for their role in promoting neural crest formation, that could explain why increasing the levels of only one of them inhibits neural crest gene expression. We therefore asked what the consequences of increased expression of both factors would be for neural crest formation. Embryos were injected in 2-cells at the 4-cell stage with either *myc* mRNA alone, *wdr5* mRNA alone (at concentrations shown to decrease neural crest factor expression), or both *myc* and *wdr5* mRNA. Strikingly, whereas each individual factor inhibited neural crest formation, co-injecting them at these same concentrations expanded the neural crest domain, as evidenced by increased expression of *foxd3* (75%, n=42), *snai2* (74%, n=50), as well as the SoxE transcription factor *sox9* (84.8%, n=33) (Fig. 4D). Together these findings support the hypothesis that wdr5 and myc work together to promote neural crest formation.

To further test this hypothesis, we asked if myc was required for the ability of low levels of wdr5 to enhance neural crest formation. Embryos were injected with myc MO, low levels of *wdr5* mRNA or both. Depletion of myc blocked neural crest formation even in embryos that expressed wdr5 at neural crest-promoting levels (*foxd3*: myc MO 83.3%, n=44; myc MO+*wdr5*: 75.6%, n=43) (Fig. 4E). These results further demonstrate that wdr5 and myc function together to positively regulate neural crest formation.

As wdr5 depletion expands the neural plate border (Fig. 2E), this suggests that endogenous wdr5 functions to restrict the size of the neural plate border. We therefore asked what effect increasing myc levels would have on this expansion. Embryos were injected with wdr5 MO, mRNA encoding myc or both and cultured to neurula stages. Increased myc activity inhibited expression of *pax3* in both the presence or absence of wdr5 depletion (*pax3*: myc only: 66.7%, n=45; myc+wdr5 MO: 85.1%, n=47) (Fig. 4F). This suggests that the relative expression levels of myc and wdr5 are crucial to proper patterning of the ectoderm.

### Mutations in wdr5 binding sites reduce binding affinity to known wdr5 binding partners

wdr5 binds to its partners through one of two conserved binding sites, the wdr5 binding motif (WBM), or the wdr5 interaction motif (WIN) (Guarnaccia & Tansey, 2018) that reside on opposite ends of the wdr5’s beta-propellor structure. The WBM site has been implicated in biding to sequence specific transcription factors such as Myc and Oct4 (Ang et al., 2011; Thomas et al., 2019) whereas the WIN site binds to arginine containing motifs (consensus ‘ARA’) present in binding partners such as KANSL1 (Dias et al., 2014) and ‘canonical’ MLL/SET domain methyltransferases, and also mediates binding to Histone H3 and chromatin (Dharmarajan et al., 2012; Patel, Dharmarajan, et al., 2008; Patel, Vought, et al., 2008; Zhang et al., 2012).

Point mutations have been identified that can disrupt the interaction between wdr5 and Myc in cancer cells (Thomas et al., 2019) (Fig 5A). To determine if these mutations can also disrupt interactions between these factors in early embryos, we used site directed mutagenesis to substitute key residues in the highly conserved WBM and WIN sites. These mutants or wildtype wdr5 were then co-expressed with myc in early embryos, and co-IP assays were carried out on blastula stage lysates. Both WBM mutants, WBM^V268E^ and WBM^N225A, L240K, V268E^, were found to reduce the interaction with myc, whereas WIN^F133A^ did not (Fig 5 B, C). By contrast, WIN^F133A^ decreased interaction with HDAC1 while mutation of the WBM site increased this interaction.

**Figure 5.**
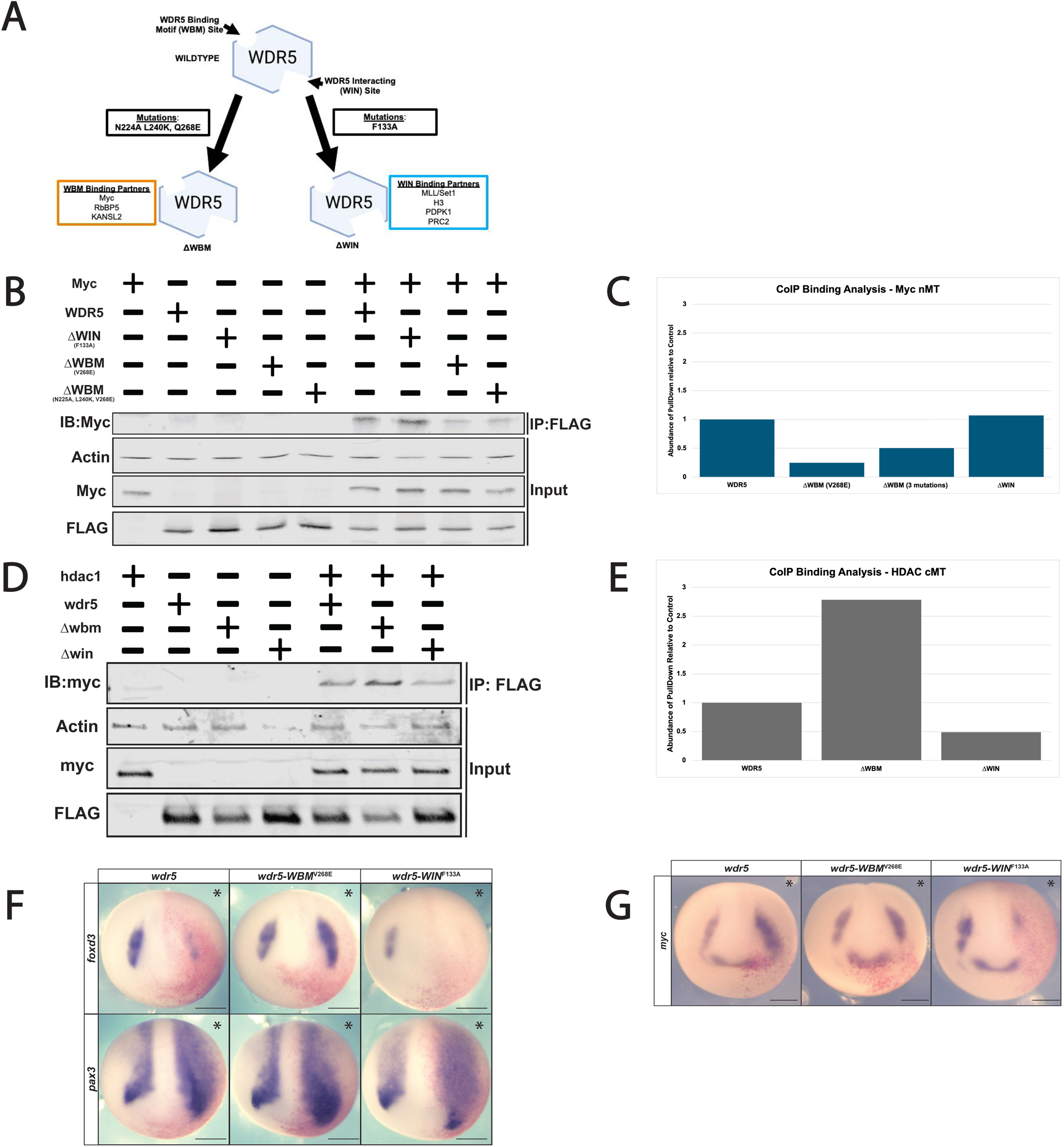
Mutations in conserved binding sites on *wdr5* differentially affect neural crest gene expression. (A) wdr5 function is primarily mediated through two-conserved and distinct binding sites, the wdr5 binding motif (WBM), and the wdr5 interacting domain (WIN), both of which can be disrupted by point mutations. (B) Western blot showcasing disruption of binding of *myc* to *wdr5-WBM^V268^ and wdr5-WBM^N225A,L240^*^K,V268E^ in *Xenopus* embryos. (C) Quantification of Co-IP binding analysis of *myc*+*wdr5, wdr5-WBM^V268^*, *wdr5-WBM^N225A,L240^*^K,V268E^, or *wdr5-WIN^F133A^* pulldown abundance. (D) Western blot showcasing disruption of binding of *hdac1* to *wdr5-WIN^F133A^* in *Xenopus* embryos. (E) Quantification of Co-IP binding analysis of *hdac1*+*wdr5, wdr5-WBM^V268^*, or *wdr5-WIN^F133A^* pulldown abundance. (F) *wdr5-WBM^V268^* exhibits an expansion of neural crest gene expression when expressed at levels that lead to loss of neural crest gene expression in *wdr5* and *wdr5-WIN^F133A^.* Overexpression of *wdr5* and all resultant mutants expand neural plate border factor expression. (G) *wdr5* mutant overexpression differentially affects myc expression compared to wildtype *wdr5*.

### Wdr5 binding mutants differentially affect regulation of the neural crest

Given that mutation of the WBM binding site decreased interaction between wdr5 and myc, and that mutation of the WIN binding site decreased interaction between wdr5 and HDAC1, we next compared the ability of these mutants to regulate neural plate border and neural crest formation as compared to wildtype wdr5. wdr5-WBM^V268E^, wdr5-WIN^F133A^ or wdr5 were expressed unilaterally at equivalent levels (Supp Fig. 4A) and injected embryos were cultured to neural plate stages for WISH. Strikingly, while wdr5-WIN^F133A^, like wildtype wdr5, led to a loss of neural crest gene expression (wdr5-WIN^F133A^ *foxd3*: 84.6% loss, n=52), wdr5-WBM^V268E^ enhanced neural crest formation (wdr5-WBM^V268E^ *foxd3*: 66% expansion, n=50) (Fig. 5F). This is likely because its reduced ability to interact with myc leads to wdr5-WBM^V268E^ functioning much like lower levels of wdr5. By contrast, expression of the neural plate border factor, *pax3*, was expanded by all three wdr5 isoforms (*pax3*: (wdr5 92.6% expansion, n=54; wdr5-WIN^F133A^ 91% expansion, n=45; wdr5-WBM^V268E^ 87.8%, n=74); further evidence that the mechanisms via which wrd5 enhances neural plate border formation are distinct from those regulating neural crest formation.

Given that wrd5 increased the neural crest domain of myc at neural plate stages, we next asked what effects the WBM and WIN domain mutants would have. Strikingly, expression of wdr5-WIN^F133A^ inhibited myc expression (*myc*: 81% loss, n=66) (Fig. 5G) unlike wildtype wdr5 (*myc*: 84.6%, n=39) or wdr5-WBM^V268E^ *(myc:* 77.2%, n=66) (Fig. 5G).

Together, these findings indicate that the WBM and WIN domains each mediate a subset of wdr5 activities with the WBM domain required for regulating definitive neural crest genes in partnership with myc. By contrast, myc itself is regulated via wdr5’s WIN-mediated functions independent of WBM– Myc binding, and this regulation appears to be less dependent upon stoichiometry.

### Both the WBM and WIN sites are required to rescue the effects of wdr5 depletion

Given the difference in phenotypes observed for the wdr5 binding domain mutants, we wished to determine the extent to which either the WBM or WIN mutant could rescue the effects of wdr5 depletion. To test this, we co-injected embryos depleted for wdr5 with mRNA encoding wdr5 at a concentration that on its own inhibits neural crest formation, or with the WBM or WIN mutants expressed at equivalent levels (Supp Fig. 5A). We observed that, although the WBM mutant on its own expands neural crest, it was unable to fully rescue neural crest formation in wdr5-depleted embryos as evidenced by lack of rescue of *foxd3* expression (*wdr5* MO+*wdr5-WBM^V268E^* mRNA: 86.5% failed to rescue, n=37) (Fig. 6A). Similarly, the WIN domain mutant was also unable to rescue wildtype wdr5 depletion (*wdr5* MO+*wdr5-WIN^F133A^* mRNA: 88.7% failed to rescue, n=53) (Fig. 6B). Taken together, the insufficiency of either mutant to rescue the loss-of-function phenotype indicates that both conserved binding sites of wdr5 are required for neural crest formation.

**Figure 6.**
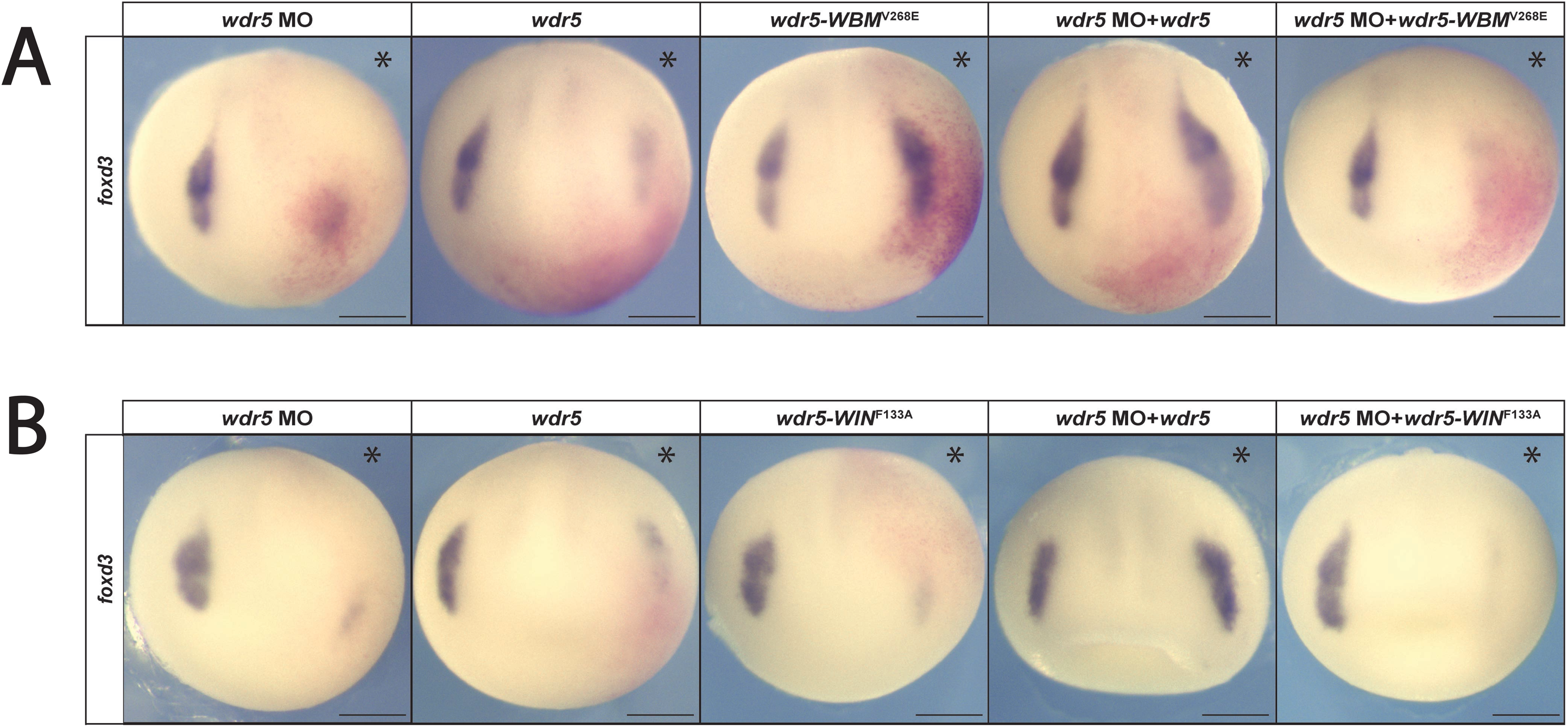
Both wdr5 binding sites are required for the expression of neural crest genes. (A) *wdr5-WBM^V268E^* overexpression fails to rescue neural crest expression in the absence of endogenous *wdr5* expression. (B) *wdr5-WIN^F133A^* overexpression fails to rescue neural crest expression in the absence of endogenous *wdr5* expression.

### Lowering myc levels promotes neural crest formation

We interpret the ability of small increases of *wdr5* or the *wdr5-WBM^V268E^* mutant to expand the neural crest domain as increasing the number of cells with the relative levels of myc and wdr5 needed to promote neural crest gene expression. If this is the case, then decreasing the levels of myc expression should have a similar effect. As 60nM of myc MO led to a complete loss of neural crest gene expression, we examined the effects that progressively lower amounts of myc MO would have on neural crest formation. Reducing the amount of MO to 30nM had little to no effect on neural crest (*snai2*: 84.4% unchanged, n=45). However, lower concentrations led to increased expression of *snai2*, consistent with our hypothesis ([15nM] 58.8% expansion, n=34; [7.5nM] 77.5%, n=40; [3nM] 85.7%, n=42) (Fig. 7A).

**Figure 7.**
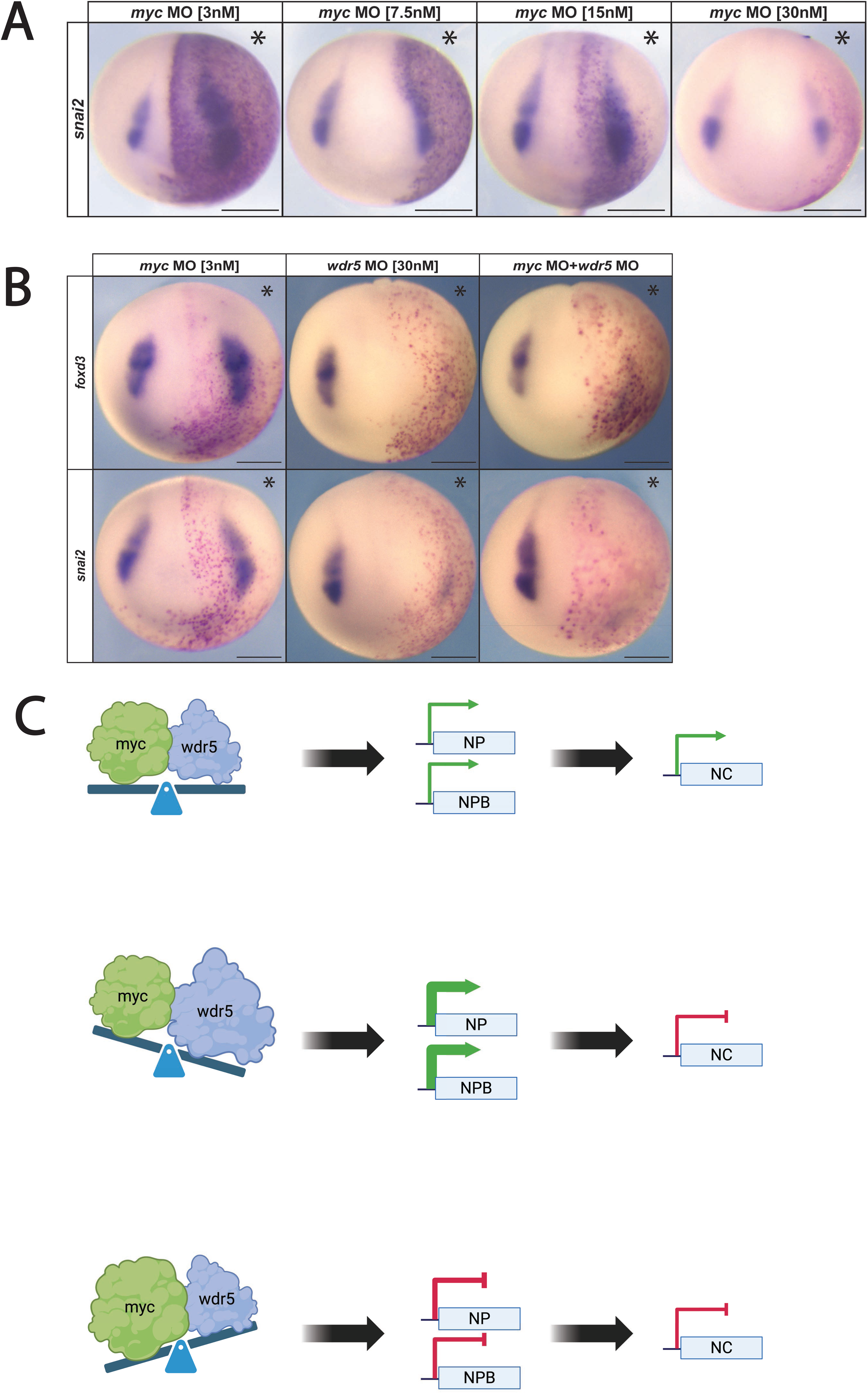
Myc downregulation expands neural crest expression via wdr5 mediated mechanism. (A) Lower concentrations of myc morpholino expand neural crest factor expression. (B) *wdr5* is required for myc-morpholino mediated expansion of neural crest gene expression. (C) Model depicting wdr5 and myc mediated regulation of the neural crest: Normal NC formation (top), loss of NC and prolonged NP and NPB (middle), loss of NC and prolonged Myc expression (bottom). Abbreviations [[NC = Neural Crest; NP = Neural Plate; NPB = Neural Plate Border]]

### Wdr5 is required for myc-mediated promotion of neural crest formation

Given that decreasing myc expression levels promotes expression of neural crest factors, we next asked whether this ability of myc to promote neural crest formation requires wdr5. Embryos were injected with myc MO at a concentration that enhances neural crest formation [3nM], wdr5 MO at a concentration that inhibits neural crest formation [30nM], or both MOs. When wdr5 was depleted, reducing myc levels was no longer able to promote expression of neural crest factors *foxd3* and *snai2*. (myc MO [3nM] + wdr5 MO [30nM]: *foxd3*: 85.3%, n=17; *snai2*: 84.6%, n=13) (Fig. 7B). This evidence supports the hypothesis that myc-mediated regulation of neural crest gene expression requires wdr5.

## Discussion

Wdr5 is a versatile scaffolding protein that functions in multiple cellular processes, including the maintenance of pluripotency in embryonic stem cells (Ang et al., 2011). Little is understood, however, about its functions in early embryonic development. Here we show that wdr5 is essential for the formation of neural crest cells in both whole embryos and induced explants, and present evidence that wdr5 plays roles in at least three distinct steps in the process via which initially pluripotent blastula stem cells transit to a neural crest stem cell state.

The most straight forward of these is in regulating the establishment of definitive neural crest cells. Here wdr5 functions with a known partner, myc, to regulate expression of genes such as *snai2* and *foxd3* in a manner dependent on its WBM domain. Intriguingly, wdr5 exhibited dose-dependent effects on neural crest gene expression. Lower doses enhanced neural crest formation, whereas higher doses inhibited it. Such biphasic effects have previously been reported for transcriptional regulators involved in pluripotency and differentiation, including Myc and Oct4 (Silva & Smith, 2008; Takahashi & Yamanaka, 2006). However, they are not characteristic of definitive neural crest factors such as snai2 or foxd3 (LaBonne and Bronner-Fraser, 2000; Heeg-Truesdell and LaBonne, 2004). Our findings suggest wdr5 levels must be precisely regulated to ensure proper stoichiometry with interacting partners such as myc. Disruption of this stoichiometry can impede neural crest formation, reinforcing the idea that developmental decisions hinge critically on balanced protein interactions.

Our data show that wdr5 and myc can physically interact in early embryos, consistent with previous studies in cancer cells (Thomas et al., 2019). similarly, our finding that co-expression of myc and wdr5 at specific doses rescued neural crest gene expression, whereas individually these factors suppressed neural crest formation, clearly highlights the functional synergy between these two proteins. Such synergy might be explained by wdr5 enhancing myc’s transcriptional regulatory functions, potentially by stabilizing complexes essential for neural crest-specific gene activation as has been observed previously (Thomas, Wang, et al., 2015). Furthermore, the finding that depletion of myc abolished the neural crest-promoting effects of wdr5 further underscores the necessity of the myc-wdr5 complex for neural crest specification.

We found that both overexpression and depletion of myc inhibited neural crest cell formation, whereas partial knockdown of myc enhanced neural crest gene expression. Similar regulatory complexity has been observed in embryonic stem cells, where partial knockdown or controlled expression of myc can promote differentiation into specific lineages (Cartwright et al., 2005). Thus, neural crest formation likely depends upon finely tuned levels of myc activity, regulated by at least in part by wdr5 interactions, to achieve proper developmental outcomes. Importantly, in cancer cells, ChIP-Seq based colocalization analyses found that approximately 80% of MYC binding sites genome-wide are also occupied by WDR5 (Thomas, Wang, et al., 2015). While disrupting the interaction between MYC and WDR5 has no effect on WDR5’s own chromatin binding, it prevents MYC from binding the majority of its target sites. The requirement for precise stoichiometry between MYC and Wdr5 has been found in both cancer cells and ES cells (Guarnaccia et al., 2021; Thomas, Foshage, et al., 2015).

Our mutational analyses also revealed a distinct role for wdr5’s WIN domain, suggesting that both differential binding partners and regulatory outcomes are controlled by these domains. Mutation of the WIN domain inhibits the expression of myc itself whereas wildtype wdr5 or the WBM mutant do not. These findings suggest that regulation of myc by wdr5 is mediated via one or more proteins that bind the WIN domain. While it could be interactions with components of chromatin remodeling complexes, it could also be a transcription factor that can interact with the WIN domain. Intriguingly, pax3 has several candidate WIN pocket interacting domains including an ARA motif at its c-terminus and WIN-binding proteins frequently present short, exposed ARA/RxR motifs near accessible, disordered tails. Investigating a role for wdr5-pax3 mediated regulation of myc will be an important future area of investigation. The results presented here show that wrd5 can mediate distinct regulatory mechanisms via separate protein domains and suggest that compartmentalization of functions within wdr5 could fine-tune transcriptional outcomes during early embryogenesis.

We also found that wdr5 depletion led to expanded expression of neural plate border markers, suggesting that it plays a role in restricting the size of the border region endogenously. Surprisingly, however, increased *wdr5* expression also promoted neural plate border formation, as did both the WBM and WIN mutants. This provides evidence that there is a third mechanism via which wdr5 regulates the process of neural crest formation that is independent of both the WBM and WIN domains. Indeed, roles for Wdr5 that do not require these domains have been previously described. For example, Wdr5 binds long noncoding RNAs (lncRNAs) via an RNA-binding pocket located between the 5th and 6th WD40 repeats, and this interaction has been shown to regulate the HoxA cluster (Lu et al., 2018; Wang et al., 2011). Furthermore, In *Drosophila*, WDR5 (WDS) binds NSL1 via a short linear motif on an outer blade surface distinct from the WIN or WBM sites (Dias et al., 2014). Future studies could profitably explore WBM/WIN-independent interacting factors responsible for neural plate border expansion. Disrupting such interaction could, for example, block cells from transiting to an epidermal state thereby retaining them in a neural plate border state.

In addition to myc, wdr5 partners with at least one other transcription factor involved in controlling pluripotency. Wdr5 has been shown to bind Oct4 in embryonic stem cells, and is required for mediating self-renewal and reprogramming via the core embryonic stem cell transcriptional network (Ang et al., 2011). Though no evidence has been shown to implicate SoxB1 factors as wdr5 binding partners, we can interpret the lack of effect on neural plate expression when *wdr5*-*WBM^V268E^* mRNA was overexpressed as an indication that interaction between wdr5 and WBM-binding partners is required for effects on *sox3* expression. It is possible that a putative wdr5 interaction with pou5f3 could explain these findings given that pou5f3 factors and soxb1 factors are required for formation of the neural plate border (Schock et al., 2024).

Given the persistent expression of both neural plate and neural plate border factors in wdr5-depleted embryos it is clear that wdr5 is not required for formation of either of these cell types, and it will be important going forward to elucidate the mechanism via which wdr5 restricts the boundaries of these domains endogenously,

Our findings are relevant to cancers such as neuroblastoma, melanoma, and pheochromocytomas that arise from neural crest derived cells (Greer et al., 1965; Guilmette & Sadow, 2019). Indeed, dysregulation of myc is estimated to underlie about a third of all cancer deaths (Tansey, 2014). The interaction between myc and the WBM binding site of wdr5 has become a key drug target in treating myc-related cancers (Thomas et al., 2019; Thomas, Foshage, et al., 2015; Thomas, Wang, et al., 2015). Likewise, the WIN binding site, which is the site that links wdr5 to chromatin and other partners, has been shown to affect a different subset of cancers when pharmacologically inhibited (Aho et al., 2019; Siladi et al., 2022). This is consistent with our findings that wdr5 can regulate multiple aspects of developmental processes in a context-dependent manner. More generally, our findings that wdr5 is essential for myc-mediated regulation of neural crest development underscores the importance of precise stoichiometric ratios of transcriptional regulators to critical developmental decisions in vertebrate embryos.

## Materials and Methods

### Animals

All animal procedures were approved by the Institutional Animal Care and Use Committee, Northwestern University, and are in accordance with the National Institutes of Health’s Guide for the Care and Use of Laboratory Animals.

### Embryological Methods

Wild-type *Xenopus* laevis embryos were staged and collected in accordance with standard methods (Zahn et al., 2022). Wild-type *Xenopus laevis* embryos were obtained using standard methods and cultured in 0.1x Marc’s Modified Ringer’s Solution (MMR) [0.1 M NaCl, 2 mM KCl, 1 mM MgSO_4_, 2 mM CaCl_2_, 5 mM HEPES (pH7.8), 0.1 mM EDTA] until the desired stages (N&F 1994). Embryos or blastula stem cell explants (also known as animal pole cell explants) used for *in situ* hybridization, HCR or immunofluorescence were fixed in 1x MEM [100 mM MOPS (pH 7.4), 2 mM EDTA, 1 mM MgSO4] with 3.7% formaldehyde and dehydrated in methanol prior to use. Embryos or blastula stem cell explants that underwent *in situ* hybridization were processed as described (LaBonne and Bronner-Fraser, 1998) and imaged using an Infinity 8-8 camera (Teledyne Lumenera). Results are representative of a minimum of three biological replicates.

Microinjection of mRNA or morpholinos was carried out at the 2-to 8-cell stage. mRNA was synthesized using an mMessage mMachine SP6 Transcription Kit (Invitrogen) and translation efficiency assessed by western blot. Either β-galactosidase mRNA or fluorescein dextran were co-injected as a lineage tracer. Approximately 10-25 ng of translation-blocking morpholinos (Gene Tools) were injected per cell. Morpholino sequences were as follows: *wdr5*: [insert sequence]; *c-myc*: [insert sequence]

### Blastula stem cell explant assays

Animal pole cells were manually dissected using forceps at blastula stage (stage 9). Manipulated embryos were injected into either both cells at the 2-cell stage or the animal cell at the 4-to 8-cell stage with either mRNA or morpholino. To induce a neural plate border/neural crest state, dissected explants were immediately cultured in 3 µM K02288 (Sigma-Aldrich) and 107 µM CHIR99021 (Sigma-Aldrich) in 1x MMR, as described by Huber and LaBonne (2024) and remained in pharmacological solution until the time of collection.

### Western blotting

Five whole embryos or ten explants were lysed in 1% NP-40 supplemented with protease inhibitors [Complete Mini, EDTA-free tablet (Roche), Leupeptin (Roche), Aprotinin (Sigma-Aldrich) and phenylmethylsulfonylfluoride (PMSF; Sigma-Aldrich)]. SDS page and western blot were used to detect proteins. The following primary antibodies were used: c-Myc 9E10(1:3000; Santa Cruz Biotechnology; sc-40); FLAG M2 (1:3000; Sigma-Aldrich; F1804); actin (1:5000; Sigma-Aldrich; A2066). IRDyes (1:20,000; mouse-800 CW; rabbit-680 TL) and the Odyssey platform (LI-COR Biosciences) were used to detect proteins. Image Studio Lite software was used to quantify protein. Results are representative of a minimum of three biological replicates.

### Co-immunoprecipitation

Five whole embryos were lysed in 1% NP-40 supplemented with protease inhibitors (see Western blotting). A 5% input was retained for western blot analysis and the remaining 95% incubated with c-Myc 9E10 antibody (1:500) for 1 hour. Approximately 25-30 µL of PAS beads (Sigma-Aldrich; P3391) were added to the lysate and incubated for 2 hours. Beads were washed with 1% NP-40 and remaining proteins eluted off the beads. Input and immunoprecipitation samples were analyzed by western blotting as described above. Results are representative of a minimum of three biological replicates.

### HCR

HCR methodologies were modified from those described by Choi et al. (2018). Whole embryos or explants were hybridized with DNA probe sets for *pax3*, *snai2* (Molecular Instruments) and incubated overnight at 37C. Probe was removed, and samples were washed and then incubated overnight with DNA hairpins labeled with Alexa 647 or Alexa 546 (Molecular Instruments). Unbound hairpins were removed by four 15-minute washes with 5x SSC and then samples were immediately mounted and imaged using a Nikon C2 upright confocal with two GaAsP detectors and four standard laser lines.

### Statistics

All statistical analyses used two-tailed *t-*tests to determine significance.

## Acknowledgements

We thank Josh York and members of the lab for thoughtful discussion. The authors acknowledge the contributions of Xenbase (https://www.xenbase.org/xenbase) and the National Xenopus Resource (https://www.mbl. edu/research/resources-research-facilities/national-xenopus-resource). Funding for the study was received National Institutes of Health (grant nos. R01GM116538 (C.L.), National Science Foundation (grant no. 1764421 to C.L.) and Simons Foundation (grant no. SFARI 597491-RWC to C.L.).

## Figure Legends

**Supplemental Figure 1.**
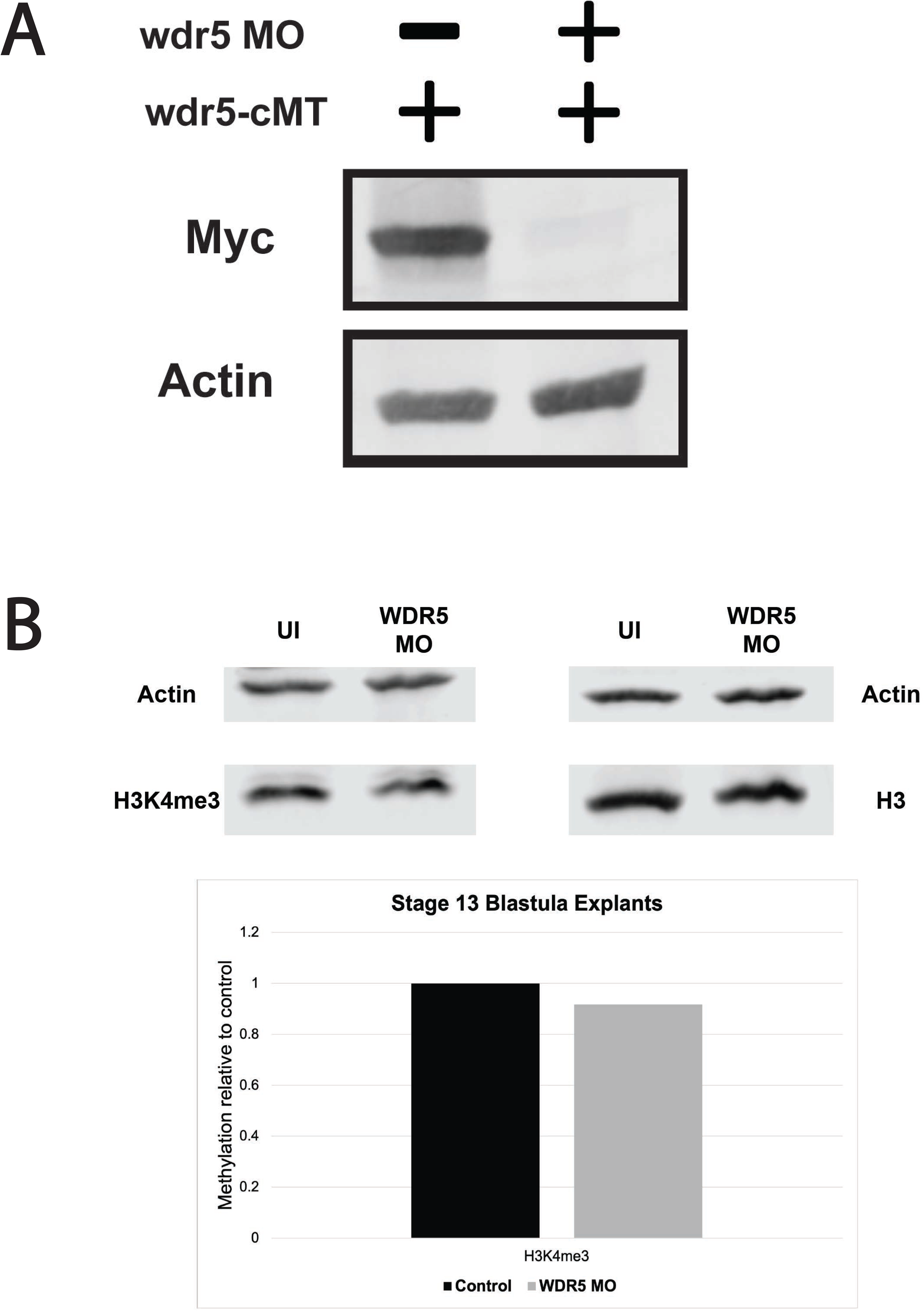
(A) Western blot depicting specificity of wdr5 morpholino. (B) Western blot and quantification of global levels of H3K4me3 compared to total H3 in control vs wdr5 MO conditions. (p>0.05)

**Supplemental Figure 2.**
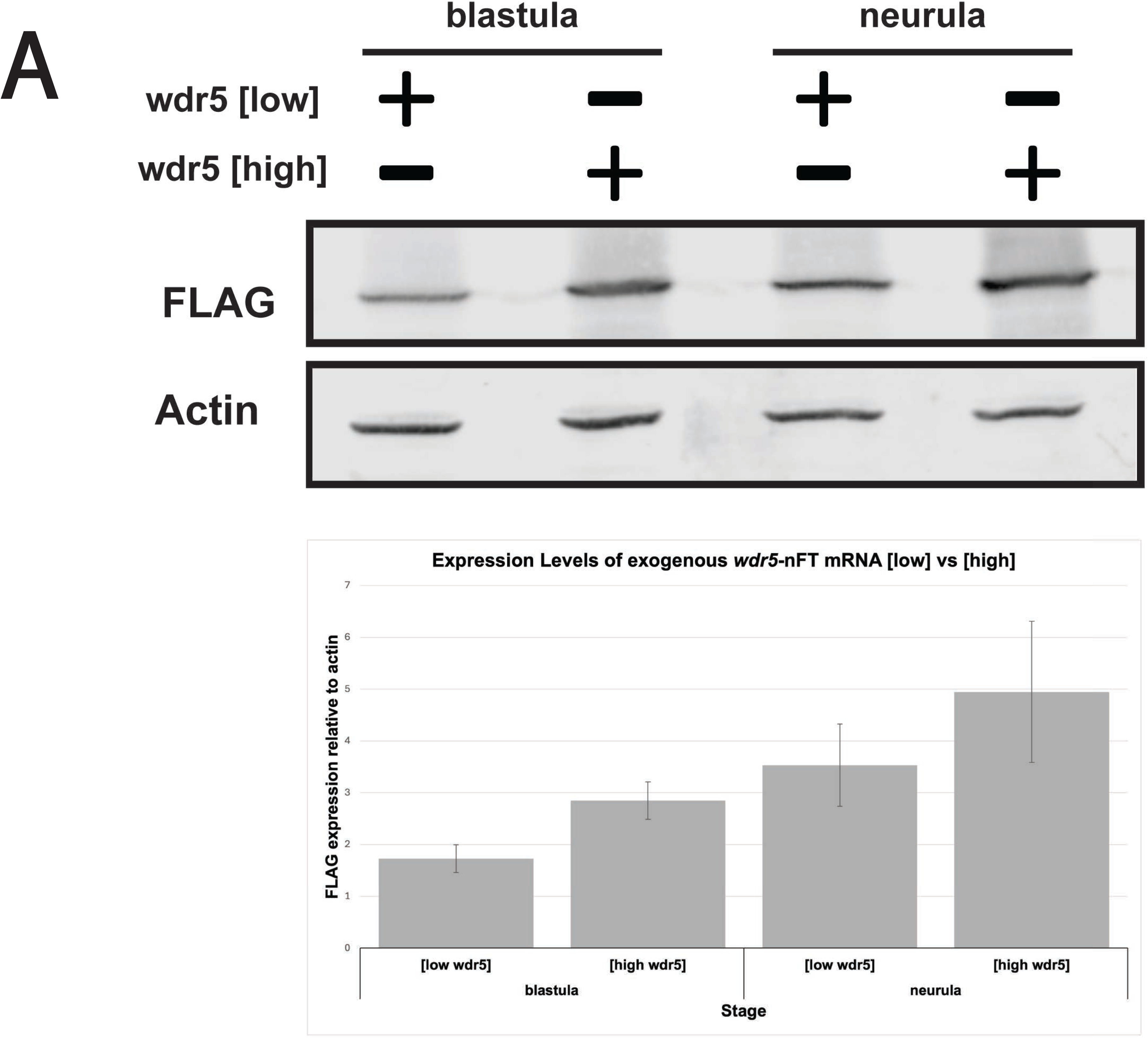
(A) Western blot and quantification of difference in low vs high concentrations of wdr*5*-nFT mRNA.

**Supplemental Figure 3.**
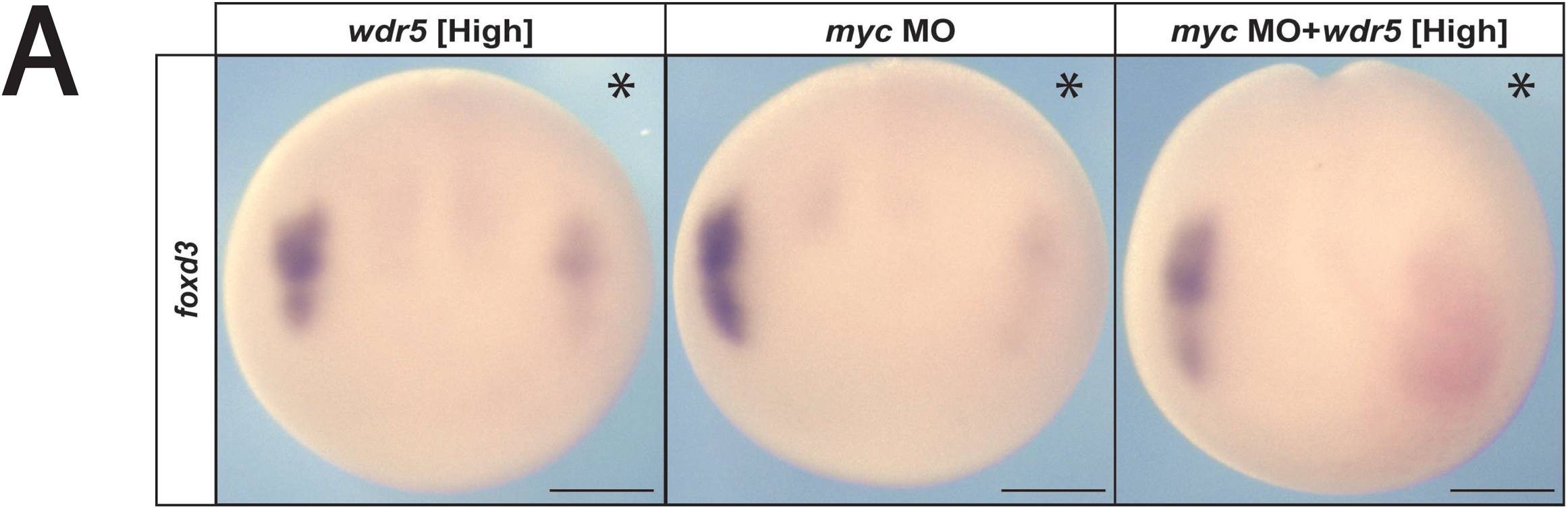
(A) High levels of wdr5 do not require myc expression to inhibit neural crest gene expression.

**Supplemental Figure 4.**
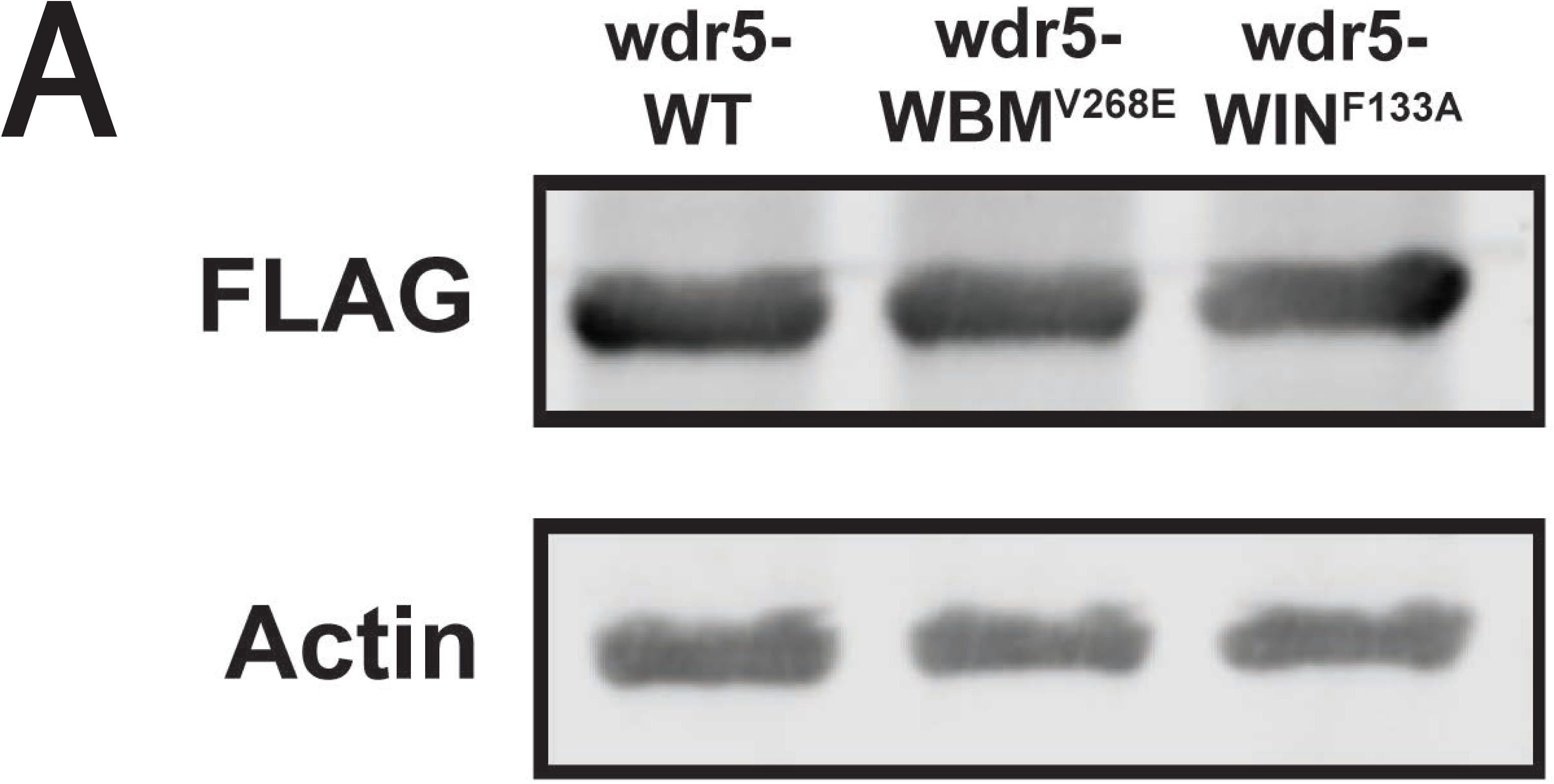
(A) Western blot showing equivalent expression of *wdr5*-*WT*, *wdr5*-*WBMV268E, wdr5*-*WINF133A* FLAG tag.

**Supplemental Figure 5.**
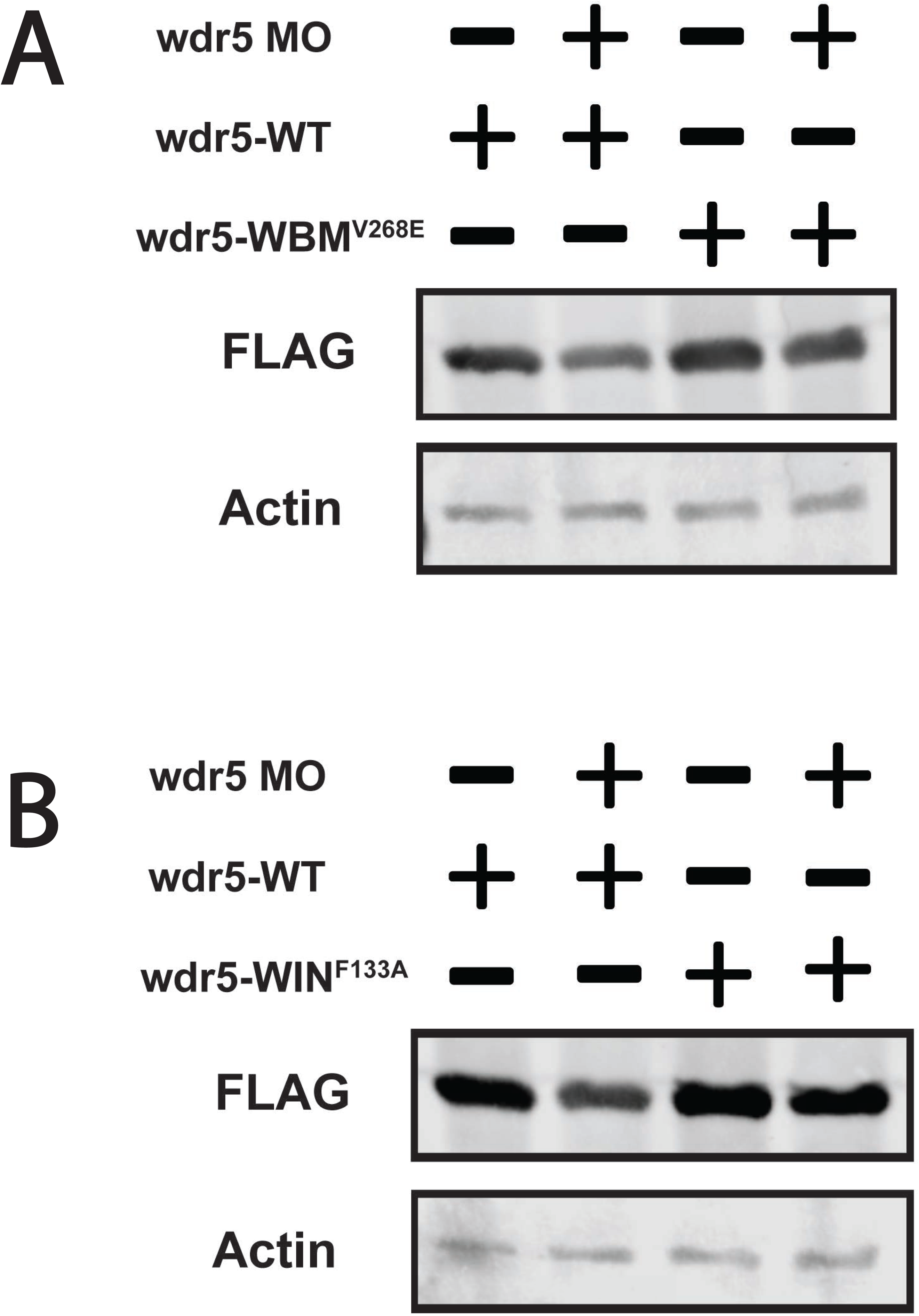
(A) Western blot showing equivalent expression of wdr5-WT and *wdr5*-*WBMV268E* in the absence of endogenous *wdr5* expression. (B) Western blot showing equivalent expression of wdr5-WT and *wdr5*-WIN*F133A* in the absence of endogenous *wdr5* expression.

